# BCG vaccination impacts the epigenetic landscape of progenitor cells in human bone marrow

**DOI:** 10.1101/2023.11.28.569076

**Authors:** Sarah J. Sun, Raúl Aguirre-Gamboa, L. Charlotte J. de Bree, Joaquin Sanz, Anne Dumaine, Leo A.B. Joosten, Maziar Divangahi, Mihai G. Netea, Luis B. Barreiro

**Author notes:** Corresponding author: Barreiro, Luis B.

## Abstract

While the Bacille-Calmette-Guérin (BCG) vaccine is used to prevent tuberculosis, it also offers protection against a diverse range of non-mycobacterial infections. However, the underlying protective mechanisms in humans are not yet fully understood. Here, we surveyed at single-cell resolution the gene expression and chromatin landscape of human bone marrow, aspirated before and 90 days after BCG vaccination or placebo administration. We show that BCG vaccination significantly alters both the gene expression and epigenetic profiles of human hematopoietic stem and progenitor cells (HSPCs). Changes in gene expression occur primarily on the most uncommitted stem cells and are reflective of a persistent myeloid bias. In contrast, BCG-induced changes in chromatin accessibility are most prevalent within differentiated progenitor cells at sites influenced by Kruppel-like factor (KLF)/SP and EGR transcription factors (TFs). These TFs are also activated in the most uncommitted stem cells, indicating that activated TFs, which drive persistent changes in HSC gene expression, likely also drive chromatin dynamics appearing within downstream progenitor cells. This perspective contests the prevailing notion that epigenetic modifications linked to innate immune memory transfer directly from stem cells to their differentiated derivatives. Finally, we show that alterations in gene expression and chromatin accessibility in HSPCs due to BCG vaccination were highly correlated (r>0.8) with the IL-1β secretion capacity of paired PBMCs upon secondary immune challenge. Overall, our findings shed light on BCG vaccination’s profound and lasting effects on HSPCs and its influence on innate immune responses.

## INTRODUCTION

The adaptive memory response is an evolutionarily conserved mechanism of vertebrate immunity. It has classically been viewed as a unique feature of T- and B-lymphocytes, which can clonally expand to generate antigen-specific memory cells with faster and more robust responses to recurrent infections. Despite this dogma, increasing evidence suggests that innate immune cells such as monocytes, dendritic cells, natural killer cells and neutrophils may harbor some antigen-agnostic memory-like properties, a phenomenon referred to as ‘innate immune memory^1,2^. This form of immune cell memory is thought to be encoded within innate immune cells through persistent epigenetic rewiring of enhancer and promoter regions of host-resistance and metabolic genes as a result of exposure to pathogens or other inflammatory signals^2^. As innate immune cells are short-lived and do not divide, it has been suggested that persistent innate immunity may be mediated by epigenetic changes within long-lived immune stem cells (hematopoietic stem and progenitor cells, HSPCs) in the bone marrow^1,2^. It has been hypothesized that these stem cells may be capable of retaining pathogen-induced epigenetic signatures across cycles of self-renewal, and transmitting these signatures to downstream innate progeny, thus maintaining a circulating pool of innate immune memory cells. While some support for this hypothesis is available from mouse models, it has yet to be fully tested in humans.

BCG vaccination is primarily used to prevent tuberculosis, but has broad ranging protective effects against a wide array of non-mycobacterial infections^3–8^. BCG vaccination protects young children in countries with high infectious pressure from all-cause mortality^9^ and infections^10^, while elderly adults receiving the BCG vaccine are significantly less likely to experience a new viral infection within the next year^6^. In murine models, BCG vaccination provides protection against heterologous (Influenza virus) infections^11^, potentially via expansion and altered gene expression within HSPCs^11,12^. Macrophages derived *in vitro* from the bone marrow of BCG-vaccinated mice demonstrate increased bacterial killing capacity for up to a year post-vaccination^12^. This suggests that gene expression changes encoded within HSPCs and passed on to developing bone marrow-derived macrophages may be responsible for some of the prolonged heterologous protective effects of BCG vaccination.

Still, the genomic mechanisms by which “memory” may be encoded in human bone marrow HSPCs, and to what extent such memory signatures are capable of rewiring the immune system’s response to infectious diseases, is incompletely understood. It was recently demonstrated that global changes in gene expression within human bone marrow persist 90 days post vaccination^13^. However, many critical questions remain unanswered, such as which HSPC cell types are most impacted, whether these expression changes are also coupled with epigenetic changes, and to what extent acquired changes in the bone marrow relate to changes in innate immune cell function.

To address these questions, we performed droplet-based scRNA-and scATAC-sequencing on the human bone marrow aspirates from 20 healthy individuals, both before and 90 days after intradermal BCG vaccination or placebo. Our data indicate that BCG vaccination impacts both the gene expression and epigenetic profiles of HSPCs for at least 90 days and that these changes are predictive of corresponding functional changes in donor-matched PBMCs challenged with *Candida albicans*.

## RESULTS

### Multimodal analysis of human bone marrow cells

Healthy individuals were randomized to receive either the intradermal BCG Bulgaria vaccine (n=15) or intradermal placebo vaccine diluent (n=5), as previously described^13^. Participants were similar in age (BCG avg = 23.7 yrs, SD 7.3, SEM 1.8; placebo avg = 21.8 yrs, SD 1.8, SEM 0.8) and sex (BCG: 5F/10M, placebo: 2F/3M). Bone marrow aspirates were collected from the iliac crest of all 20 individuals prior to vaccine administration (D0), and 90 days after vaccination (D90) and cryopreserved for future processing (Fig 1A). To isolate HSPCs from each bone marrow sample, we stained bone marrow aspirates with fluorescence-conjugated antibodies targeting CD34, a transmembrane phosphoglycoprotein specific to HSPCs^14^. We also stained all bone marrow aspirates with a panel of antibodies targeting canonical immune cell markers (CD3, CD56, CD14, etc. for mature immune cells and CD90, CD10, CD110, etc. to distinguish between CD34+ HSPC subtypes, Table 1, Fig 1B). We used fluorescence activated cell sorting to sort out live, CD34+ HSPCs for downstream droplet-based scRNA-seq processing (115,698 cells captured) and scATAC-seq (58,988 cells captured) processing while simultaneously collecting flow cytometry data. This workflow enabled the simultaneous collection of single cell gene expression, chromatin accessibility, and surface protein data for each sample (Fig 1A).

**Figure 1.**
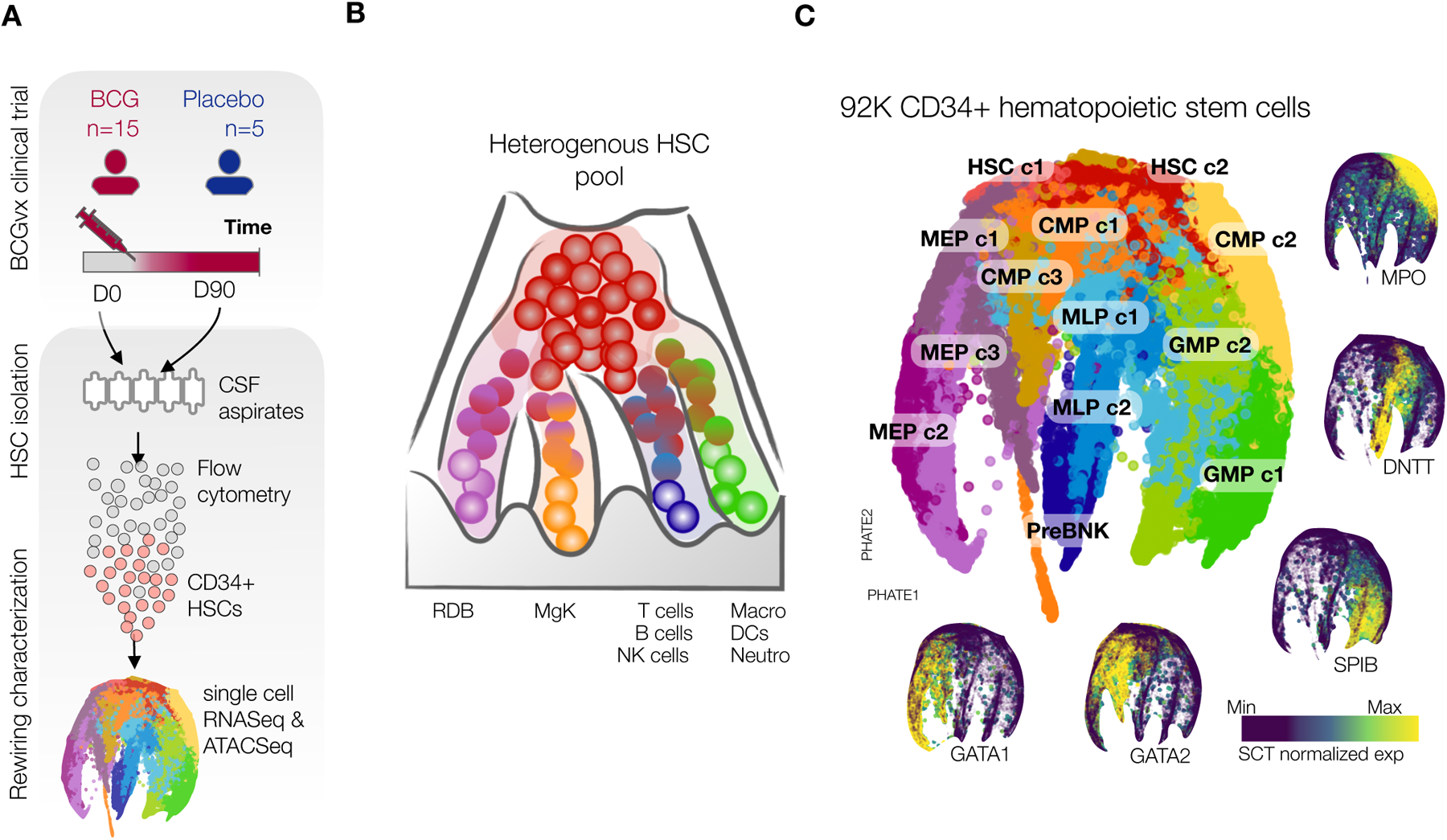
Multimodal analysis of human bone marrow cells. **(A)** Overview schematic of experimental timeline and samples collected in a clinical trial setting for BCG vaccination, 20 total donors on BCG (n=15) or placebo (n=5). Bone marrow aspirates and PBMCs were collected from all patients at D0 prior to vaccination and D90 (90 days after BCG or placebo). Cryopreserved bone marrow samples were stained with a cocktail of lineage and HSPC-specific antibodies to enable flow cytometric analysis of cellular composition as well as simultaneous sorting of all CD34+ cells. Sorted CD34+ cells were immediately processed for scRNA-seq and scATAC-seq according to the respective 10X genomics protocols. **(B)** Revised model of hematopoiesis in which the developmental process is a continuum, as recent findings indicate that only a small fraction of HSC generate an equal outcome for all blood mature cell lines, while most HSC exhibit a differentiation bias toward one lineage. **(C)** PHATE of the scRNA-seq data collected from bone marrow CD34+ HSPCs of BCG and placebo vaccinated individuals at D0 and D90. Cells were grouped into 13 non-overlapping clusters based on gene expression: HSC c1 (n=9637), HSC c2 (n=10953), CMP c1 (n=9174), CMP c2 (n=14918), CMP c3 (n=1715), GMP c1 (n=6631), GMP c2 (n=6423), MEP c1 (n=8871), MEP c2 (n=5439), MEP c3 (n=3811), MLP c1 (n=5153), MLP c2 (n=3837), PreBNK (n=3371). Marker gene colored PHATE plots by expression levels of lineage defining genes (from left to right) GATA1, GATA2, SPIB, DNTT, MPO.

### BCG vaccination has a long-term impact on gene expression within HSPC populations

Given that BCG vaccination was previously shown to impact gene expression in bulk RNA-seq of HSPCs^13^, we first asked whether the 90-day impact of BCG vaccination on gene expression within HSPCs was detectable at single-cell resolution, and what cell types within HSPCs were the most responsive to BCG vaccination. To assess the effect of BCG vaccination on gene expression of different HSPC subtypes, we clustered all high-quality cells in the scRNA-seq dataset into 13 non-overlapping groups and then assigned them to known HSPC cell subtypes (HSCs, CMPs, MLPs, GMPs, MEPs, and Pre-BNK cells) (Fig 1C), based on the expression of pre-determined lineage-specific genes such as GATA1 (erythroid), DNTT (lymphoid), MPO (myeloid/neutrophilic), and SPIB (pDC) (Fig 1C).

To investigate the overall HSPC signature after BCG vaccination, we collapsed the single-cell gene expression values for each of the 13 main clusters to generate pseudobulk estimates for each sample. Then, we used a mixed linear model to identify genes for which expression levels changed in response to BCG vaccination, while controlling for temporal changes independent of BCG vaccination (as measured in the placebo samples, Fig S1A, B). We used a multivariate adaptive shrinkage model (mash^15^), which leverages the correlation structure of effect sizes across cell types to increase power and refine effect size estimates. Genes with a stringent local false sign rate (lfsr) < 0.01 were considered *differentially regulated (DR)* due to BCG vaccination (Fig 2A,B, Table 2).

**Figure 2.**
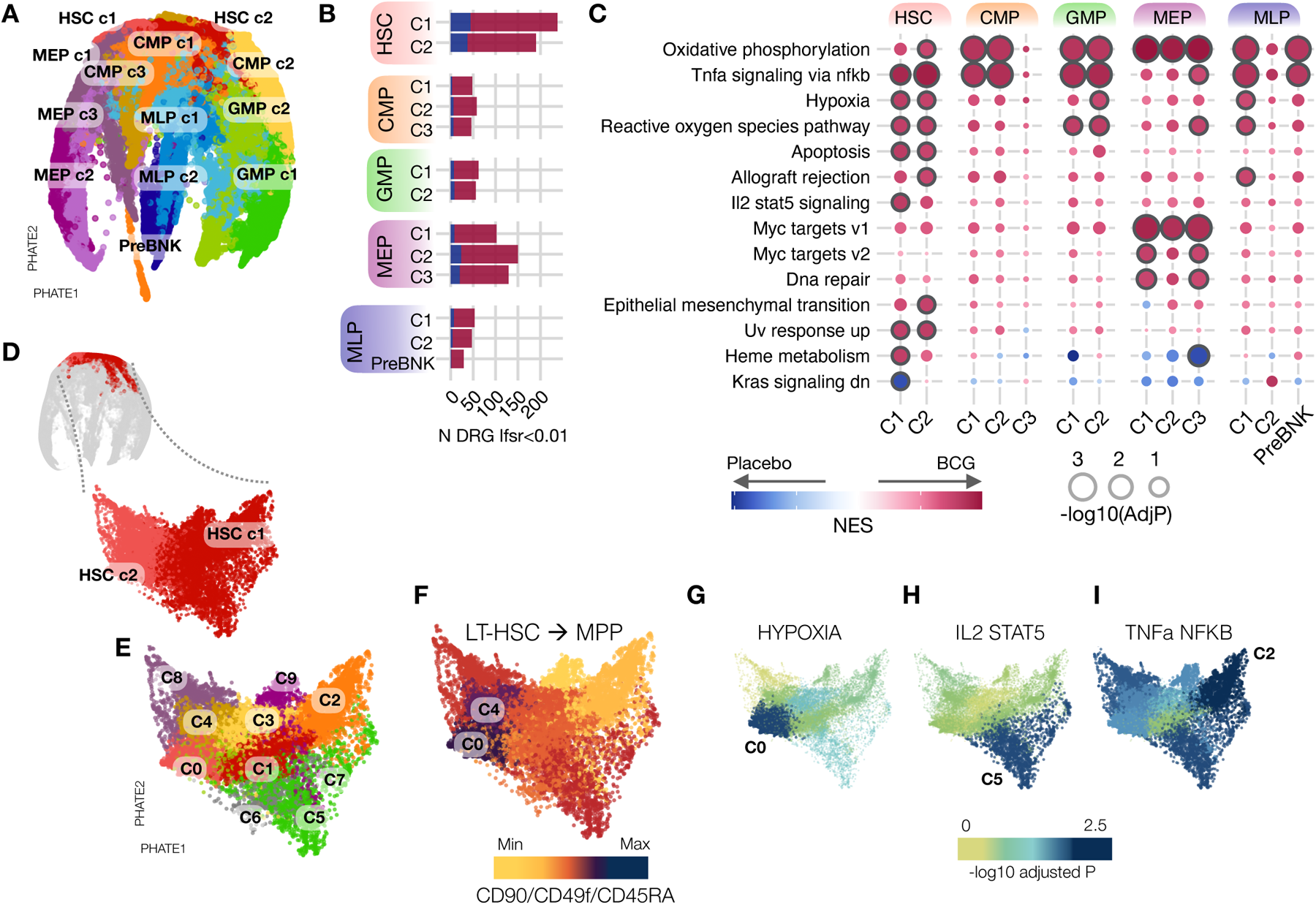
BCG vaccination has a long-term impact on gene expression within HSPC populations. **(A)** PHATE of the scRNA-seq data collected from bone marrow CD34+ HSPCs of BCG and placebo vaccinated individuals at D0 and D90. **(B)** Bar graphs summarizing the total number of significant genes (lfsr<0.01) in each cell type. Blue and red shading indicate the number of genes whose expression was impacted negatively and positively, respectively, by BCG vaccination compared to placebo. **(C)** Summary plot of gene set enrichment analysis (GSEA) performed separately for each cell type. Genes were ordered by the rank statistic – log10(pval)*logFC and compared against Hallmark gene sets. Circle size is scaled to – log10(padj). All shown circles are pathways with p<=0.05. All circles with border have padj<=0.1 **(D)** Second-round clustering was performed on HSC c1 and HSC c2 **(E)** HSC subgroups after second round clustering of HSC c1 and HSC c2 cells. **(F)** HSC subclusters colored by CD90/CD49f/CD45RA (average z-score across the three genes). Darker shading indicates a higher score. **(G-I)**. UMAPs of select Hallmark pathways. Shading of each subcluster is scaled to –log10(padj) enrichment of the pathway within the cluster.

DR genes were heterogenous across HSPC cell types, with the largest percentage residing within the stem-cell clusters HSC c1 and HSC c2 (Fig 2B). Strikingly, in HSC c1 and HSC c2, 238 and 190 genes respectively were differentially regulated (lfsr<0.01) 90 days following BCG vaccination, with 59.7% of total DR genes in HSC c1 and c2 shared between the two clusters. Megakaryocyte-erythroid progenitor (MEP) clusters had the second highest number of DR genes (MEP c1 = 102, MEP c2 = 150, MEP c3 = 129). In contrast, no other cluster had more than 62 DR genes, demonstrating that a single intradermal BCG vaccination preferentially impacts the gene expression landscape of the most long-term, self-renewing stem cells in the bone marrow at three months following vaccination.

Upon bacterial infection or LPS challenge, cytokines such as IL6, IL3, IL1β, G-CSF, TNF, IFNγ, and GM-CSF are produced either by non-hematopoietic cells or by HSPCs themselves^16–18^. However, our data showed that their gene expression levels were either very low (median log counts per million reads < 1) across all cell types (Figure S1C) or showed no significant difference between BCG and placebo groups (with a padj > 0.9). This suggests that by day 90, it is unlikely that HSCs or other cells are actively responding to any remaining vaccine antigens or pathogen-associated molecular patterns (PAMPs).

DR genes were enriched for immune, metabolism, or proliferation/apoptosis pathways (Fig 2C, Table 3). Some immune-related pathways such as IL2/Stat5 signaling were enriched (padj < 0.1) predominantly within HSC clusters, suggesting that immune rewiring was greatest within the most undifferentiated stem cells. Others were more ubiquitous, such as ‘TNF via NFκB signaling pathway’ and ‘oxidative phosphorylation’ which were enriched across the spectrum of HSPC subtypes, representing a subset of universally modulated pathways. MEPs exhibited a unique metabolic signature compared to all other HSPC subtypes, with predominant enrichments in MYC signaling (critical in regulation of cellular metabolism, proliferation, differentiation, and apoptosis) but smaller enrichments in stress-related pathways (hypoxia, reactive oxygen species) that were more predominant in HSC and other progenitor clusters. Principal component analysis comparing HSPC subtypes based on gene set enrichment scores revealed a tight and distinct clustering of HSC c1 with HSC c2 and of MEP c1 with MEP c2 and MEP c3, away from the zero-reference point (Fig S1D). HSCs and MEPs clustered on opposite ends of PC1, further demonstrating the strong but differential impact of BCG vaccination on these cell types. Overall, the data show that BCG vaccination impacts the expression of immune, metabolism, and proliferative genes across HSPC subtypes for at least 3 months, but that the stem-like HSCs and progenitor MEPs have the strongest propensity to maintain a lasting state of differential gene expression. Enrichments in TNF signaling and oxidative phosphorylation were uniquely pervasive across all cell types (Fig 2C), leading us to speculate that these gene expression differences reflect the select retention of specific immune and metabolic gene expression programs sustained within HSCs as they differentiate into downstream progeny.

### BCG vaccination heterogeneously impacts HSCs

Our gene expression data indicated that HSC c1 and HSC c2 are the primary sites of differential gene expression 90 days after BCG vaccination. We reasoned that changes within these clusters may be of particular significance, given that HSC c1 and HSC c2 are capable of giving rise to all downstream progenitors. Thus, we decided to delve deeper into understanding how these cell populations were modified. In order to investigate BCG-induced gene expression changes within HSCs at higher resolution, we performed a refined clustering on cells classified as HSC c1 or HSC c2 into 10 total HSC subgroups (Fig 2D, E). We classified each subcluster as a Long Term(LT)-HSCs, Short Term(ST)-HSC, or multipotent progenitor (MPP) based on its expression of markers *CD90*, *CD49f*, and *CD45RA,* which have classically been used to differentiate between these HSC subtypes^19^ (Fig 2F). As we had previously done for each major cluster, we performed a differential gene expression analysis and gene set enrichment analysis for each HSC subcluster. Enrichments of pathways that were significant within HSC c1 and c2 (Fig 2C, Table 3) were variable when assessed indivudally within each HSC subcluster (Fig S1E). Many pathways had the strongest enrichment within phenotypically LT-HSC – like subsets such as c0 and c5. Interestingly, pathways enriched predominantly within HSCs (Fig 2C), but not within downstream progenitors were almost exclusively enriched only within LT-HSC-or ST-HSC-like clusters, but not MPPs. For example, the HSC-specific ‘Hypoxia’ pathway was enriched only within the most phenotypically LT-HSC subcluster c0 (Fig 2G; p.adj = 0.0039) and the pathway ‘IL2-Stat5 signaling’ was enriched only within a phenotypically ST-HSC subcluster c5 (Fig 2H; p.adj = 0.0048). In contrast, the ‘TNF via NFκB signaling pathway’ which was significantly enriched not only within HSCs, but also within the most downstream progenitor clusters (CMP, GMP, MEP, MLP, and PreBNK), had the strongest enrichment in the phenotypically MPP-like subcluster c2 (Fig 2I, p.adj = 0.0018). Our data indicate that gene expression changes within HSCs appear to represent the heterogenous rewiring of LT-HSCs, ST-HSCs, and MPPs. Moreover, gene expression changes within MPP-like HSCs may be associated with transmission to downstream progeny, while gene expression changes harbored by the more quiescent LT-HSCs and ST-HSCs are less likely to be transmitted or shared by downstream cell types.

### BCG vaccination results in lineage bias of HSCs

Severe bacterial infections can induce a state of emergency myelopoiesis, in which the bone marrow increases the production of myeloid cells which circulate in the bloodstream and extravasate into sites of infection^16^. Well-controlled, localized infections (such as BCG vaccination) are not typically associated with the induction of emergency myelopoiesis. Yet, numerous studies in mice have suggested that exposure to BCG can rewire the bone marrow towards myelopoiesis^20,21^, at least acutely. Whether this is true in humans and could represent a persistently rewired state rather than an acute reaction, has not been investigated. Given that the detection of persistent gene expression changes was within HSCs, we next investigated whether the altered baseline of immune and metabolic gene expression programs in HSCs of BCG-vaccinated individuals was associated with a persistent skewing of HSC lineage bias (Fig 3A,B).

**Figure 3.**
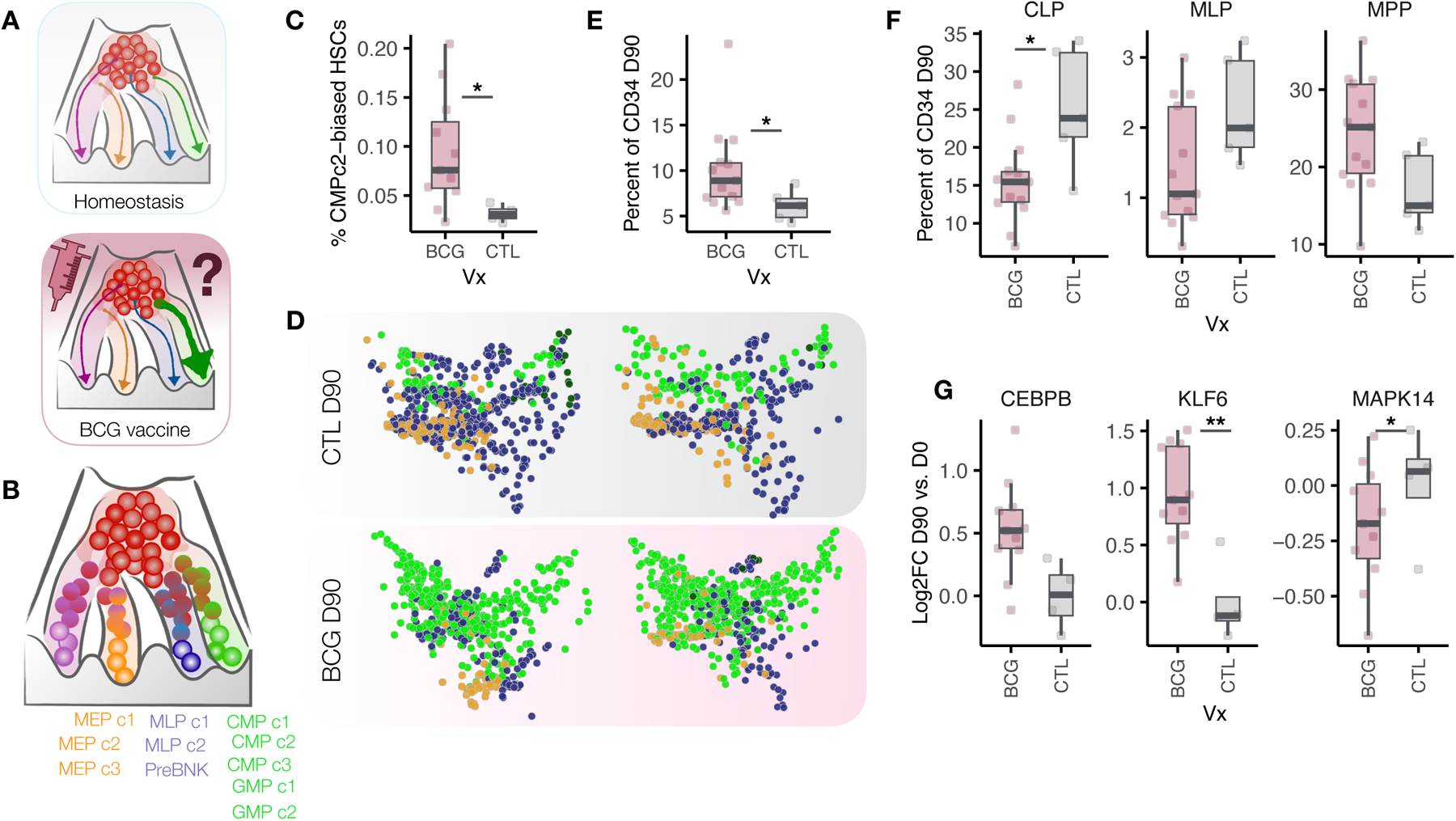
BCG vaccination results in lineage bias of HSCs. **(A)** Schematic of the experimental question. **(B)** Model of hematopoiesis showing possible terminal fates used in the Cellrank model. **(C)** Absolute percentages of CMP c2 biased HSCs for each donor at D90 (p=0.018). **(D)** Terminal fates of individual HSCs from BCG and placebo vaccinated individuals were predicted with CellRank for each donor. PHATE maps show HSCs from placebo individuals (top) or BCG-vaccinated individuals (bottom) colored by predicted terminal fate (green: CMPs/GMPs, blue: MLPs/PreBNK, orange: MEPs). **(E)** Percent CMPs in the bone marrow of 90 days post BCG (pink) compared to placebo (blue), Mann Whitney p = 0.026. **(F)** Bar graphs showing the percentage of each cell type among live CD34+ HSPCs at D90 as determined by flow cytometry analysis. CLP p = 0.026, MLP p = 0.075, MPP p = 0.095. **(G)** D90 vs. D0 Log2FC expression of *CEBPB* (lfsr = 0.1), *KLF6* (lfsr = 0.001), *MAPK14* (lfsr = 0.03)

To make lineage bias predictions within single cells we utilized CellRank^22^, a similarity-based trajectory inference method that utilizes RNA-velocity information (measurements of unspliced to spliced mRNA) to infer developmental directionality of HSCs at single-cell resolution within a snapshot in time (at D90).

Comparing the composition of lineage biases between BCG and placebo individuals at day 90, we found that BCG vaccination led to a significant increase in the percentage of HSCs biased towards the CMP c2 terminal state (p = 0.018, Fig 3C, D). In support of this prediction, flow cytometry analyses of the bone marrow aspirates revealed an overall increase in CMPs among BCG-vaccinated relative to placebo individuals (Fig 3E; p=0.026), accompanied by a decrease in the percentages of CLP and MLP (Fig 3F) among BCG vaccinated individuals. Assessment of marker genes specific to CMP c2 revealed these cells to be the predominant expressors of *CEBPB* (Figure S2A), *CSF3R* (Figure S2B), *MPO* (Figure S2C), and *CEBPA* (Figure S2D), establishing their likely identity as neutrophil or *MPO*-expressing monocyte progenitors.

BCG-vaccination increased the gene expression of *CEBPB* (lfsr = 0.1) within HSC c1, as well as decreased expression of *MAPK14* (lfsr = 0.03), a prototypical p38 MAPK, which can act as an inhibitor of granulopoiesis^23^. Other transcription factors involved in inflammatory responses, such as *KLF6* (lfsr = 0.001), which interacts with *NFkB* to promote pro-inflammatory gene expression within mature myeloid cells^24–26^, and *IRF1* (lfsr = 0.004) which coordinates proinflammatory gene expression in response to viruses and bacteria^27,28^, were also significantly increased in HSC c1 upon BCG vaccination (Fig 3G, S2E). Together, these data show that BCG vaccination has a long-term impact both on genes involved in inflammation within HSCs and genes that promote neutrophil/monocyte bias and differentiation. This results in HSCs which harbor a combination of altered baseline gene expression programs and lineage biases towards myelopoiesis.

### BCG vaccination impacts the chromatin accessibility of immune progenitors

Epigenetic alterations are believed to be central to innate immune memory. Thus, we next investigated whether the gene expression signatures detected in our scRNA-seq data were associated with changes in the epigenetic landscape of HSPCs. To investigate the epigenetic effects of BCG we used sample-paired scATAC-sequencing data collected on CD34+ HSPCs before and after BCG vaccination (Fig 1A). After quality-control filtering, 58,988 high-quality cells were retained (see methods). These cells were clustered into 16 cell populations (Fig 4A). Initial cell type annotations (HSC, CMP, GMP, MEP, MLP, and PreBNK) were generated by mapping chromatin accessibility-based ‘gene activity’ scores calculated for well-established lineage-specific genes (Fig 4B) to gene expression within the scRNA-seq data (Fig 1C), and transferring scRNA-seq labels to the closest matched clusters in the scATAC-seq data. scATAC-seq clusters mapped to the same broad cell type were then given a subtype label (HSC 1, 2, etc). In the same way we had previously detected *differentially regulated* genes, we asked whether BCG vaccination led to changes in peak accessibility (*differentially accessible*, or DA, peaks). We identified more than 13,000 total DA peaks (Figure 4C, Table 4) across all clusters, demonstrating that BCG vaccination not only had lasting impacts on gene expression, but also on the epigenetic landscape of HSPCs. The largest total number of these DA peaks were located within peripheral myeloid CMP, GMP, and MEP clusters, while fewer DA peaks were found within HSCs, despite these being the cell types harboring the most gene expression changes.

**Figure 4.**
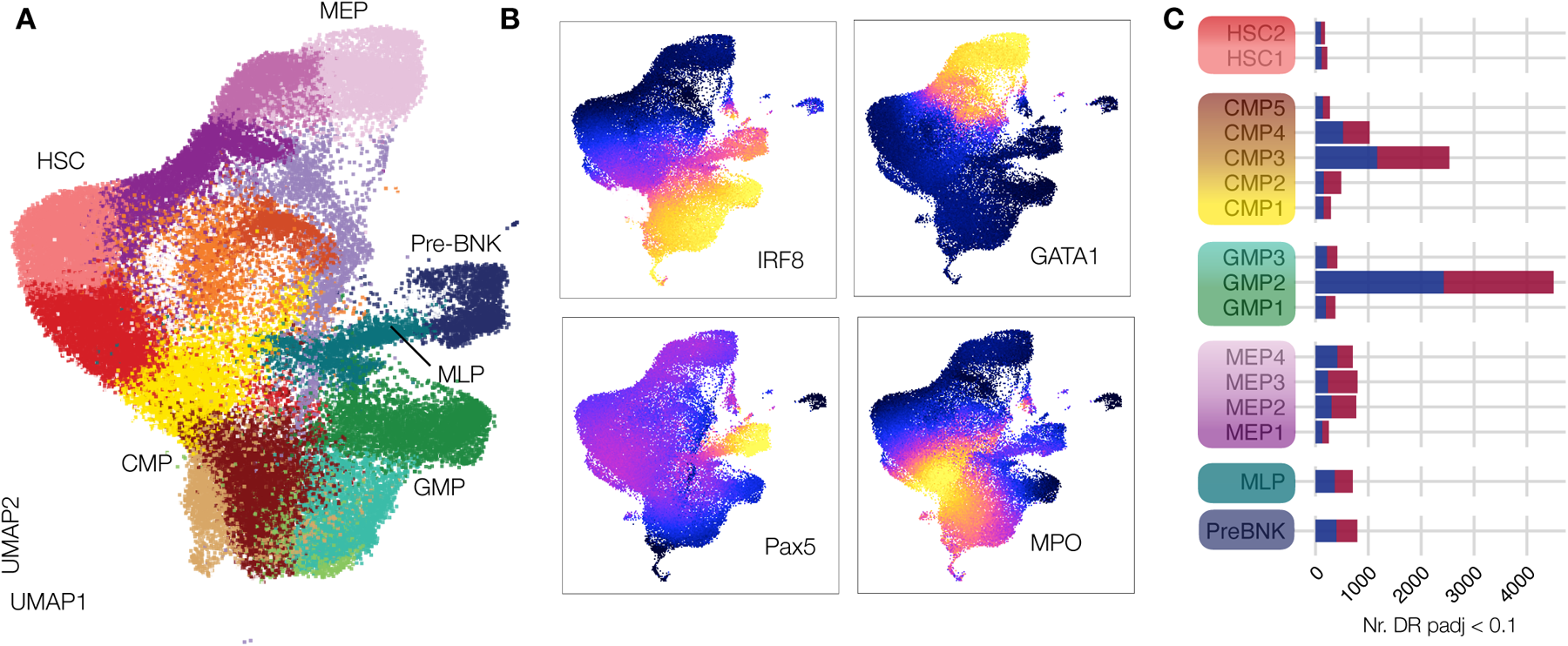
BCG vaccination impacts the chromatin accessibility of immune progenitors. **(A)** UMAP of the scATAC-seq data collected from bone marrow CD34+ HSPCs of BCG or placebo vaccinated individuals at D0 or D90. Clustering based on chromatin accessibility grouped cells into 16 clusters: CMP1 (n=5041), CMP2 (n=2942), HSC1 (n=5750), HSC2 (n=5052), MEP1 (n=4436), PreBNK (n=1876), GMP1 (n=3798), CLP (n=2146), MEP2 (n=2653), MEP3 (n=3886), CMP3 (n=1458), MEP4 (n=2230), CMP4 (n=1952), CMP5 (n=5590), GMP2 (n=962), GMP3 (n=2806). **(B)** UMAPs colored by gene activity scores of lineage-defining genes (IRF8 – DC; GATA1 – MEP; PAX5 – lymphoid; MPO – granulocytic/myeloid). Gene scores are indicative of the degree of chromatin accessibility within a 100 kb window on either side of the gene body (dark = lo, bright yellow = hi). **(C)** The total number of significant peaks (FDR<0.1) for each cluster. Blue and red bars indicate peaks whose accessibility was impacted negatively and positively, respectively, by BCG vaccination compared to placebo.

Clusters CMP3 and GMP2 harbored the greatest changes in chromatin accessibility, although many DA peaks were also detected within MEPs and other CMP/GMP clusters. Next, we aimed to identity what TFs are most likely responsible for the observed changes in chromatin accessibility. To do so, we combined related clusters into six main cell type groups (HSC, CMP, GMP, MEP, MLP, and PreBNK), and then searched for TF motifs that were enriched within the combined DA peaks of each cell type (Fig 5A). Not surprisingly, the strongest enrichments resided within CMPs and GMPs and included TFs families such as EGR1-4, NRF1, and E2F6 with significant roles in macrophage immune modulation, cholesterol metabolism, and DNA methylation respectively (Table 5). The top three motif classes within CMPs were KLF/SP1 (fdr = 10^-66^), TCFL5 (fdr = 10^-45^), and E2F6 (fdr = 10^-41^). The top motifs within GMPs were highly overlapping, including the top three motifs, KLF/SP1 (fdr = 10^-36^), E2F6 (fdr = 10^-22^), and EGR1-4 (fdr = 10^-19^). When excluding motifs with low abundance (present within > 15% DA peaks) top enrichemnts within both CMP and GMP included the motif families EGR1-4, CTCF, and KLF/SP1 (Fig 5A).

**Figure 5.**
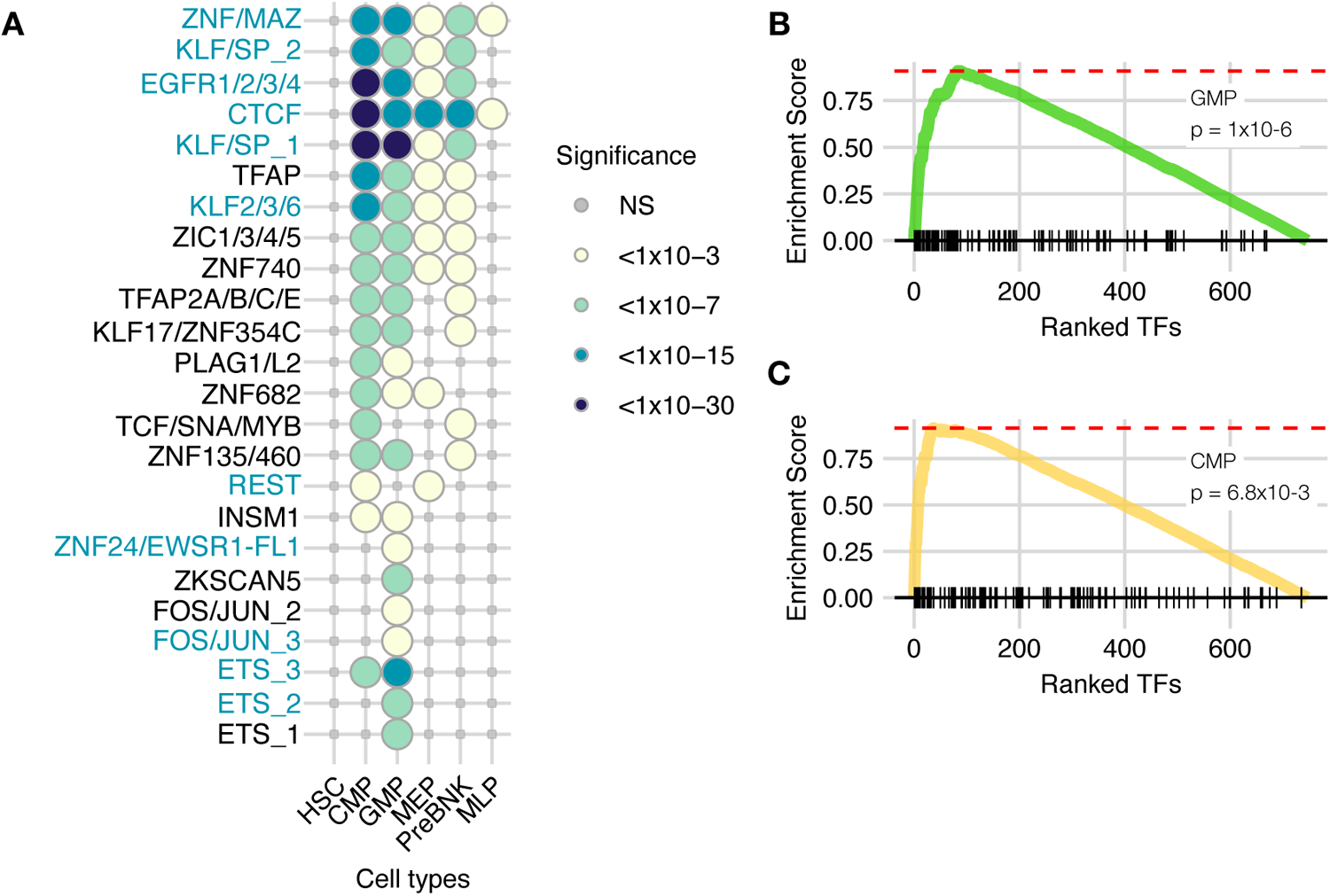
Changes in chromatin accessibility in downstream myeloid progenitors are coupled to TF-driven rewiring of HSCs. **(A)** Top transcription factor motif enrichments for broad cluster groups. Circle color is scaled to –log10(FDR). TFs shown with a circle have FDR < 0.001 and are present in at least 15% of DR peaks in at least one cell type. Areas with no circle indicate an enrichment with FDR >= 0.001. TFs highlighted in teal color also serve as gene expression drivers within HSCs **(B-C)**. GSEA enrichment plots showing the enrichment of DA-peak associated motifs of GMP **(B)** and CMP **(C)** within TFs that exhibit differential activity in HSCs following BCG vaccination.

We then asked whether these TFs were the same TFs responsible for driving the differential gene expression programs detected within HSCs upstream. To this end we used the differential gene expression data from our scRNA-seq analysis (Fig 2) to make inferences about the differential activity of TFs underlying gene expression changes within HSCs. We performed a TF activity analysis utilizing the SCENIC^29,30^ pipeline, using our single cell gene expression data to search for evidence of up- or down-regulation of transcription factor *modules* within HSCs, each defined by a central driving TF and all of its predicted target genes. In this analysis, the differential expression of many gene targets of a TF is interpreted as evidence of altered TF activity. Despite limited power, we detected several TF modules with evidence of differential activity (p < 0.05) induced by BCG within HSCs and MEPs (Fig S3A,B, Table 6). Strikingly, we found that the TF motifs enriched among DA regions in CMP and GMP are significantly enriched within TFs that exhibit differential activity in HSCs following BCG vaccination (Fig 5B-C, p = 1x10^-6^ and 6.8x10^-3^ in GMP and CMP, respectively). Overall, our data suggest a model wherein BCG-vaccination modulates the activity of a unique subset of TFs to induce a prolonged low-level myeloid bias and differential gene expression program within HSCs, that results in the establishment of a changed epigenetic landscape of downstream myeloid clusters.

### BCG-induced differential chromatin accessibility within myeloid-like HSPCs predicts increased IL1**β** secretion by PBMCs

Given that CMPs and GMPs retained extensive epigenetic signatures, we asked whether chromatin accessibility changes within myeloid progenitors could have functional implications for the mature immune cells they give rise to. When assigned to their closest genes, DA peaks in CMPs and GMPs are enriched for reactome pathways related to TLR2 and TLR4 signaling and signaling by interleukins (Fig 6A), as well as biological process pathways such as ‘neutrophil activation’ and ‘neutrophil degranulation’ (Figure S4A, Table 7), which was the strongest within CMPs. DA peaks within other HSPCs (MEP, MLP, and PreBNK) had minimal enrichments, in agreement with lower numbers of DA peaks within those cell types.

**Figure 6.**
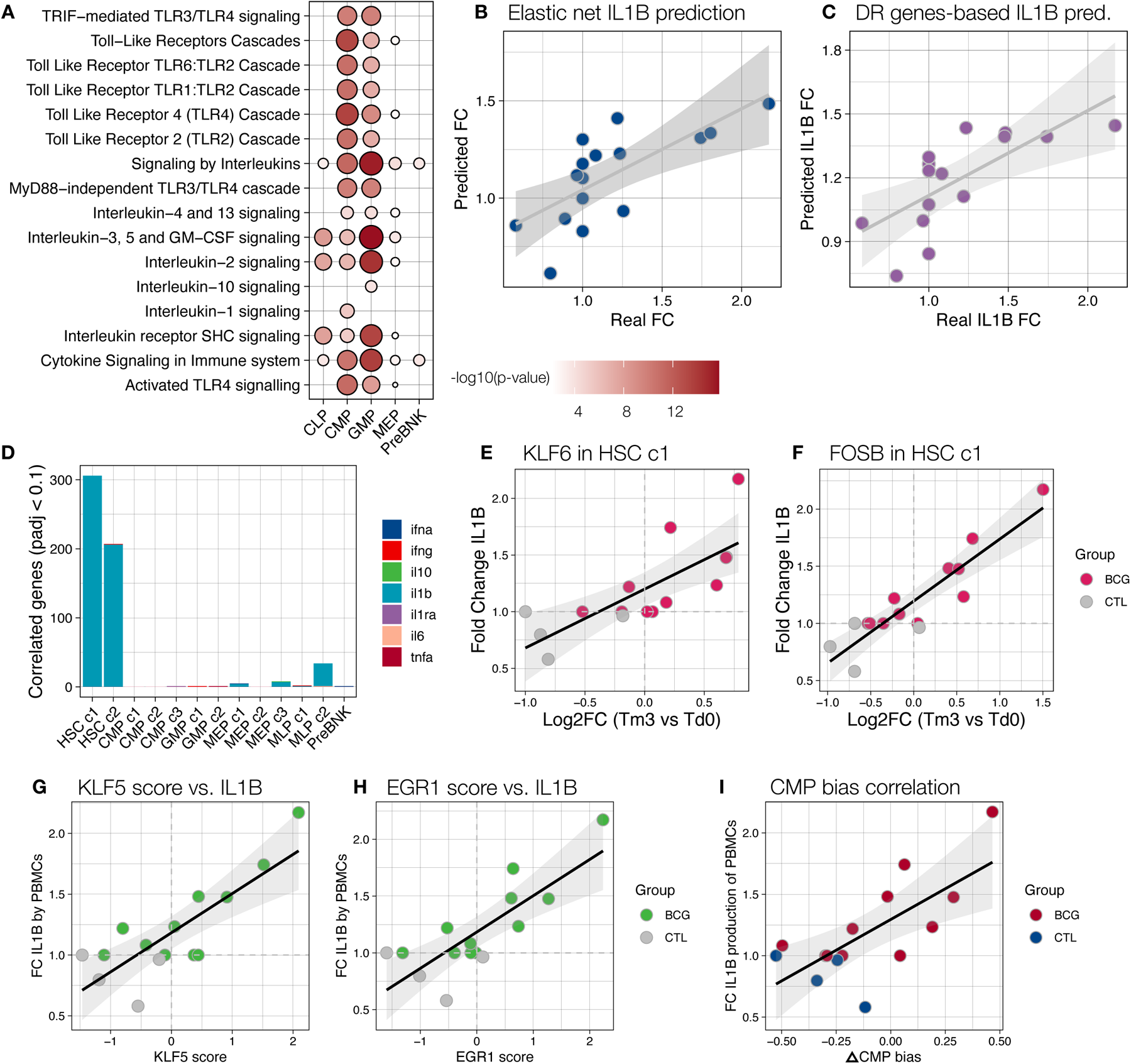
BCG-induced differential chromatin accessibility within myeloid-like HSPCs predicts increased IL1β secretion by PBMCs. **(A)** DA peaks within each cluster were assigned to the gene with the closest TSS. Gene ontology enrichment analysis was performed for reactome pathways using all peak associated genes as background and genes associated with DR peaks as foreground. Plot circle size and shading darkness are both scaled to –log10(p-value) of enrichment. **(B)** Results of the elastic net regression. The scatterplot shows real IL1B FC (D90 vs. D0) for each donor on the x-axis and predicted values from the model trained using DA peaks in CMP5 on the y-axis (Spearman rho = 0.761, p = 0.001). **(C)** Results of the elastic net regression using gene expression data for HSCs. The scatterplot shows real IL1B FC (D90 vs. D0) for each donor on the x-axis and predicted values from the model on the y-axis (Spearman rho = 0.837, p = 1x10-4). **(D)** Quantification of the total number of DR genes in HSCs with significant spearman correlations with fold change IL1B secretion for each cytokine tested. **(E-F)** Spearman correlations between log2FC expression levels of the transcription factors KLF6 (Spearman rho = 0.86) and FOS (Spearman rho = 0.83) and fold change IL1B secretion between D0 (before) and D90 (3 months-post BCG vaccination). **(G-H)** Example scatter plots correlating TF activity (regulon) scores in HSCs with fold change IL1B in PBMCs for KLF5 (spearman rho = 0.71, p = 0.0033) and EGR1 (spearman rho = 0.71, p = 0.0029). **(I)** Plot correlating the fold change IL1B secretion between D0 (before) and D90 in PBMCs with the △CMP lineage bias (percent HSCs biased towards CMPs (CMP c1, c2, or c3) at D90 vs. D0) for each individual (Spearman rho = 0.63, p = 0.01).

These results suggested that DA peaks within myeloid progenitors could regulate innate immune pathways, and that CMP- or GMP-derived innate immune cells entering the peripheral blood could have altered immune functionality. As a measure of peripheral blood cell immune functionality, we used previously collected cytokine secretion data from donor-matched PBMCs^43^ generated by Cirovic *et al*^13^. The data contain a panel of secreted cytokine concentrations by the PBMCs of each donor in response to a 24-hour stimulation with heat killed *C. albicans,* and showed that BCG, compared to placebo vaccination, rewired PBMCs to produce increased levels of IL1β and IL6 in response to *C. albicans* challenge^13^. Although the BCG vaccination cohort generally secreted higher amounts of these proinflammatory cytokines, individual PBMC samples displayed a high degree of intra-cohort heterogeneity, suggesting that the effects of BCG vaccination likely rewired PBMC cytokines responses to varying degrees across individuals. We reasoned that if differential accessibility at DA peaks were directly involved in the reprogramming of these cytokine responses, one would expect to find individuals with the greatest magnitude of epigenetic rewiring to be the same individuals with the greatest increases in cytokine secretion. To test this hypothesis, we used elastic net regression to formally determine whether levels of differential accessibility at DA peaks (raw D90 vs. D0 log2FC values) had power to predict cytokine responses (FC D90 vs. D0) across individual donors. This demonstrated that the fold change increase in IL1β production could be predicted by log2FC DA peak accessibility to a high level of accuracy (Figure 6B; R = 0.761, p = 0.001) within the peripheral MPO^hi^ CMP cluster, CMP5, but not other clusters. These data suggest that changes in chromatin accessibility that have a meaningful impact on IL1β cytokine secretion are specific to a single lineage and cell type.

We asked whether the same individuals harboring the largest changes in CMP5 chromatin accessibility and increased cytokine secretion were the same individuals harboring the greatest BCG-induced changes in the activity of driver transcription factors and differential gene expression in HSCs. As done using DA peaks, we used elastic net regression to determine whether levels of differential expression of DR genes within HSCs had power to predict IL1β responses (FC D90 vs. D0) across individual donors (Figure 6C). Differential gene expression within HSCs had strong and significant predictive power (R = 0.837, p=1x10^-4^), establishing that changes in IL1B secretion capacity are also tightly linked to day-90 differential gene expression within HSCs. Log2FC responses of hundreds of individuals genes within HSCs correlated significantly (padj < 0.1) with IL1β responses, further supporting the elastic net regression findings (Figure 6D). Importantly, the log2FC values of several transcription factors such as KLF6 (Figure 6E; in the KLF2/3/6 family) and FOSB (Figure 6F) were among these significantly correlated genes with R > 0.8. Moreover, the *activity* scores of several key transcription factors (including EGR1 and KLF5 whose motifs were strongly enriched within CMP DA peaks) within HSC had remarkably strong correlations with IL1β production (Figure 6G-H, S4B-G). Finally, we found a significant correlation between IL1β production and the extent of BCG-induced CMP bias within HSCs (Fig 6I) as determined using cellrank (Fig 3). Collectively these data formally demonstrate that BCG-induced differential TF activity and gene expression in HSCs, induction of lineage bias towards CMP, downstream CMP progenitor chromatin accessibility, and peripheral immune cell cytokine secretion are linked processes.

## DISCUSSION

Since clinical evidence and mouse models have suggested that the BCG vaccine may impact the immune system at the hematopoietic stem-cell level^12,31–33^, we used single-cell RNA and ATAC sequencing on HSPCs isolated from human bone marrow aspirates to investigate how BCG vaccination affects gene expression and chromatin accessibility 3 months later. While the effect of BCG vaccination on innate and adaptive immunity have been studied extensively using samples from peripheral blood, our study is the first to assess the impact of BCG vaccination on gene expression and chromatin accessibility at single cell resolution, using samples derived directly from healthy adult human bone marrow. Here we found that BCG vaccination alters the programming of gene expression, TF acitivty, lineage bias, and chromatin accessibility in a celltype specific manner within bone marrow cells. These features are variably modifed across individuals, and have significant power to predict IL1β secretion from donor paired PBMCs in response to a heterologous *C. albicans* challenge. These data support the hypothesis that long-lasting activation within uncommitted HSCs directly influences the epigenetic landscape of downstream progenitors, which enter the circulation as functionally reprogrammed cells (Fig 7). Our model suggests that HSCs residing at the top of the differentiation hierarchy act as the central drivers of all changes detected downstream. Thus, our data raise critical questions about the drivers that maintain the re-wired TF circuits and gene expression programs within HSCs. Do small numbers of bacteria persist three months following BCG vaccination? Or did the initial exposure to BCG induce a self-sustained, persistent epigenetic rewiring of HSCs? In support of the latter possibility, BCG was not detected in microbiological and molecular tests of the bone marrow samples^13^, suggesting that bacteria are cleared within three months of vaccination, or otherwise persist at undetectable levels. Likewise, HSPCs from BCG vaccinated individuals did not secrete increased levels of pro-inflammatory cytokines which direct immune activation of HSCs is known to induce. Instead, our data suggest that the baseline expression program within HSCs is altered by transient exposure to BCG, at least for intermediate (90 day) time scales. Future epigenetic analyses aimed at characterizing DNA methylation levels, for example, within HSCs could help determine the molecular drivers maintaining differential gene expression programs within HSCs.

**Figure 7.**
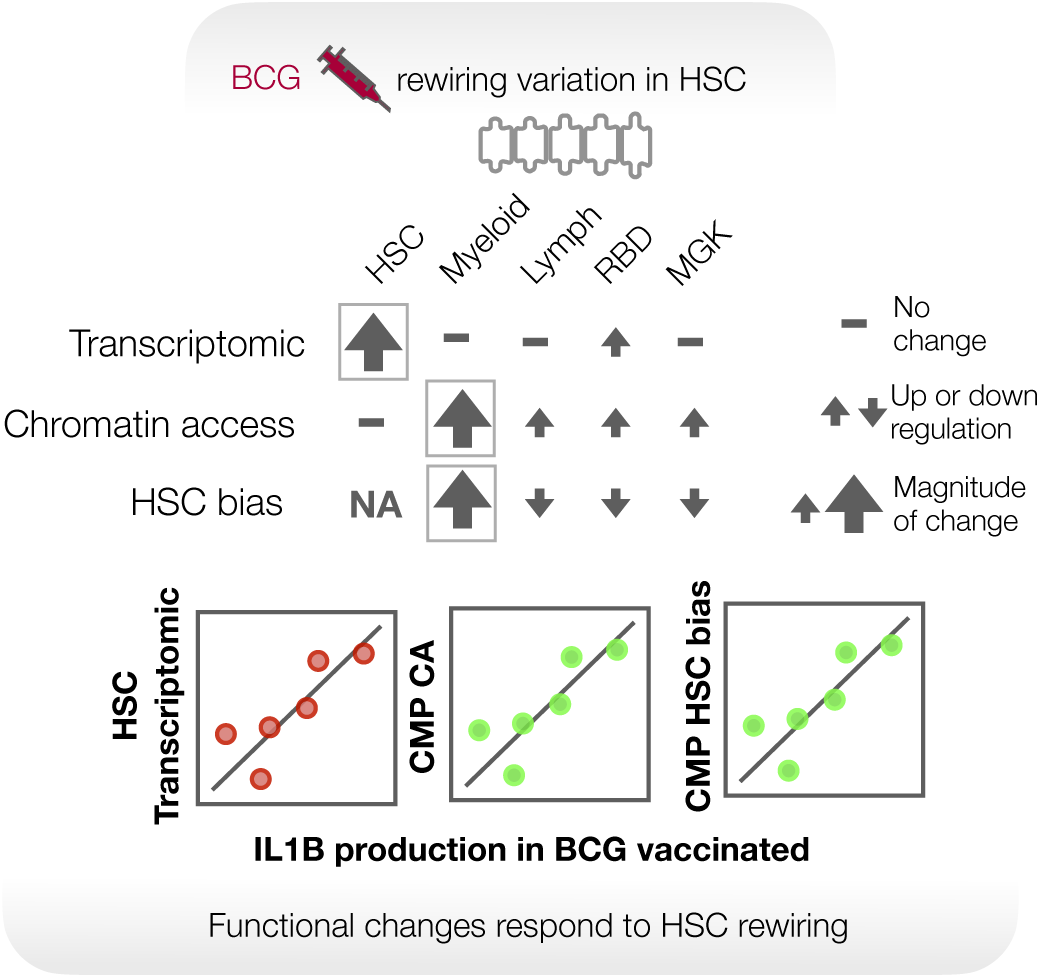
Summary Figure. BCG vaccination rewires TF activity and gene expression within HSCs for at least 3 months. Progenitors deriving from these HSCs harbor changes in chromatin accessibility at thousands of sites. These sites are predicted to serve as binding sites for TFs with altered activity within HSCs. Multiple features within the bone marrow, including the magnitude of differential gene expression change within HSCs, TF activity, myeloid bias, and chromatin accessibility changes within CMPs are predictive of changes in IL1B cytokine section by donor-paired PBMCs.

Interestingly, the gene expression and chromatin accessibility signatures of BCG vaccination appear to affect different populations of progenitor cells. As emphasized above, HSCs harbored rewired myeloid-driving gene expression programs driven by altered transcription factor activities, while the resulting downstream myeloid progenitors harbored changes in chromatin accessibility, but not large numbers of DR genes. We hypothesize that these changes in chromatin accessibility serve as an epigenetic memory signature of altered transcription factor activities and binding patterns occurring at the HSC-stage of development that is subsequently imparted onto the downstream progeny, even as many of these differential gene expression programs are silenced. When performing formal comparisons of baseline (in D0 samples) peak accessibility between HSC and CMP, we have found that consistently higher percentages of peaks (∼25%) have decreased (log2FC < 0; P.adj < 0.1), compared to increased (∼11%) accessibility in CMP compared to HSC. Likewise, a peak in GMP is more likely to have decreased accessibility compared to the same peak within HSC (33.6% with significant reduction in accessibility compared to 17.7% of peaks with a significant increase), suggesting that quantitative losses in peak accessibility are a general feature of hematopoietic differentiation. It is likely that, as HSCs differentiate into the myeloid lineage, the gene expression programs regulated by these now closed regions of accessibility are also lost, allowing myeloid progenitors to retain only select differential gene expression programs such as those affecting TNF, NFκB, and oxidative phosphorylation programs. The differential accessibility landscape within these progenitors then may largely reflect the incomplete closing of peaks around the binding sites of TFs that were differentially active within upstream HSCs, leading to memory-like signatures. Future work involving direct investigations into TF binding through ChIP-like approaches within HSCs and the myeloid progenitors derived from them will be critical for further investigating this model.

Finally, this study represents the first single cell analysis to directly show that immune induced changes to both chromatin accessibility and differential gene expression patterns within human bone marrow are directly predictive of functional responses in the periphery. These findings demonstrate the systemic nature by which vaccines impact the immune system and highlight the fact that human vaccination with live attenuated vaccines can have lasting impacts not only on adaptive lymphocytes, but also on central compartments such as the bone marrow for at least three months. Ultimately, our data suggest that the impact of BCG on the bone marrow is very complex due to 1) high levels of HSPC heterogeneity, including at baseline conditions, and 2) the dynamic nature of continuous cell differentiation that continues to take place even as HSPCs are responding to the vaccine. We suggest future work consider lineage tracing to directly examine the inheritance of gene regulatory marks from stem to progenitor cells. More broadly, we hope that more work will focus on understanding the durability of central stem cell reprogramming, the types of pathogens against which this phenomenon might be protective, the inter-individual factors that dictate differences in HSPC responsiveness to BCG vaccination between individuals, and how, or whether, the unique sequences of infectious and immune challenges encountered by humans over the course of a lifetime may imprint unique HSPC memory fingerprints reflected within the innate immune cell compartment.

## EXPERIMENTAL MODEL AND SUBJECT DETAILS

### Bone marrow aspirate staining, sorting, and sample collection

The study enrolled twenty BCG-naive volunteers, both male and female, aged between 18 and 50 years, as described in^13^. These participants were healthy, with no active infections, no signs of inflammation, and a negative Quantiferon-TB Gold test result. They were randomly divided into two groups: 15 participants were administered a standard dose (0.1ml) of intradermal BCG vaccine (BCG Bulgaria, Intervax), while the remaining 5 received 0.1ml of a placebo vaccine diluent. The trial was approved by the Arnhem-Nijmegen Ethical Committee (approval number NL55825.091.15). Cryopreserved bone marrow aspirates were processed, following the steps detailed below, on 7 separate days/batches, each batch containing 1) males and females and 2) samples collected on both D0 and D90. Five out of the seven batches contained samples from both placebo and BCG vaccinated cohorts (two contained only BCG cohort samples when there were no remaining controls).

#### Initial thawing and incubation

Cryopreserved samples were thawed and cultured in RPMI 1640 (Fisher) supplemented with 10% fetal bovine serum (Corning), 2 mM L-glutamine (Fisher), 2% HEPES (Thermo Fisher Scientific), 1% non-essential amino acids (Thermo Fisher Scientific), 1% essential amino acids (Thermo Fisher Scientific), 0.14% 5N NaOH, 1mM sodium pyruvate (Thermo Fisher Scientific), 100U/ml penicillin (Thermo Fisher Scientific), and 100μg/ml streptomycin (Thermo Fisher Scientific) for 2 hours. After incubation, samples were washed with PBS, passed through a 100 µm filter, and counted.

#### Antibody staining

To prepare samples for flow cytometry analysis and sorting, cells were incubated with 1:50 Live/dead fixable blue (Invitrogen) at a final cell concentration of 1M cells/100 µL for 20 mins (on ice). Samples were washed with 1% BSA (Miltenyi Biotec) in PBS (used for all further washing and staining steps) and resuspended in F/C block solution (BD Biosciences) for 10 minutes. Cells were washed and resuspended in a cocktail of antibodies targeting mature and stem/progenitor cell surface markers (See Table 1) for a final cell concentration of 1M cells per 100 µL. After 30 minutes on ice, cells were washed, resuspended, and passed through a 70 um Flowmi cell strainer (Fisher) immediately prior to sorting.

#### Sorting

Sorting was performed on a Symphony S6 cell sorter in the UChicago Human Disease and Immune Discovery core (HDID) using a 100 µm nozzle. For any given batch, samples collected from the same donor were sorted sequentially alternating between starting timepoints (for example, batch1: S1 D0, S1 D90, S2 D0, S2 D90; batch 2: S3 D90, S3 D0, S4 D90, S4 D0). Following sorting, CD34+ cells were washed in 1% BSA in PBS, counted, and then processed for single cell RNA and ATAC captures are described below:

#### Single cell RNA capture

Immediately prior to capture, samples were combined into two pools (2 or 3 samples per pool). Multiplexed cell pools were used as input for the single cell captures. For pools containing 2 or 3 samples, 6600 cells or 10,000 cells respectively were targeted for collection using the Chromium Single Cell 3’ Reagent (v3.1 chemistry) kit (10X Genomics). Post Gel Bead-in-Emulsion (GEM) generation, the reverse transcription (RT) reaction was performed in a thermal cycler as described (53°C for 45 min, 85°C for 5 min), and post-RT products were stored at -20°C until downstream processing.

#### Single cell ATAC capture

Leftover cells in each pool not used for single cell RNA capture were lysed for 3 minutes to isolate nuclei, transposed, and used as input for the single cell ATAC captures. Variable numbers of nuclei (ranging from 2,026 to 9,085, depending on the number of leftover cells) were targeted for collection using the Chromium Next GEM Single Cell ATAC Reagent (v1.1 chemistry) kit (10X Genomics). Post Gel Bead-in-Emulsion (GEM) generation, the GEMs were incubated in a thermal cycler as described (72°C for 5 min, 98°C for 30 sec, 12 cycles of 98°C for 10 sec, 59°C for 30 sec and then 72°C for 1 min), and post-incubation products were stored at -20°C until downstream processing.

#### Bulk CD34-processing

Total RNA was extracted from the sorted CD34-cell fraction of each sample using the miRNeasy Micro kit (Qiagen) or miRNeasy Mini kit (Qiagen). RNA-sequencing libraries were prepared using the Illumina TruSeq protocol. Indexed cDNA libraries were pooled in equimolar amounts and sequenced single-end 100 bp reads on an Illumina NovaSeq.

### Single cell library preparation and sequencing

#### Single cell RNA libraries

Post-RT reaction cleanup, cDNA amplification, and sequencing library preparation were performed as described in the Single Cell 3’ Reagent Kits v3.1 User Guide (10X Genomics). Briefly, cDNA was cleaned with DynaBeads MyOne SILANE beads (ThermoFisher Scientific) and amplified in a thermal cycler using the following program: 98°C for 3 min, 11 cycles x 98°C for 15 s, 63°C for 20 s, 72°C for 1 min, and 72°C 1 min. After cleanup with the SPRIselect reagent kit (Beckman Coulter), the libraries were constructed by performing the following steps: fragmentation, end-repair, A-tailing, SPRIselect cleanup, adaptor ligation, SPRIselect cleanup, sample index PCR (98°C for 45 s, between 11 and 13 cycles x 98°C for 20 s, 54°C for 30 s, 72°C for 20 s, and 72°C 1 min), and SPRIselect size selection. Prior to sequencing, all multiplexed single-cell libraries were quantified using the KAPA Library Quantification Kit for Illumina Platforms (Roche) and pooled in an equimolar ratio. Libraries were sequenced 100 base pair (read1: 28, i7: 10, i5: 10, read2: 90) on an Illumina NovaSeq.

#### Single cell ATAC libraries

Post GEM incubation cleanup and sequencing library preparation were performed as described in the Single Cell ATAC Reagent Kits v1.1 User Guide (10X Genomics). Briefly, post-incubation GEMs were cleaned up first with DynaBeads MyOne SILANE beads (ThermoFisher Scientific) and then with SPRIselect reagent (Beckman Coulter). Libraries were constructed by performing sample index PCR (98°C for 45 s, 9 or 10 cycles of 98°C for 20 s, 67°C for 30 s, 72°C for 20 s, and 72°C 1 min) followed by SPRIselect size selection. Prior to sequencing, all multiplexed single-cell libraries were quantified using the KAPA Library Quantification Kit for Illumina Platforms (Roche) and pooled in an equimolar ratio. Libraries were sequenced 100 base pair (read1: 50, i7: 8, i5: 16, read2: 50) on an Illumina NovaSeq.

## QUANTIFICATION AND STATISTICAL ANALYSIS

### Mapping, demultiplexing, and cell filtering

#### Single-cell RNA-seq data

FASTQ files from each multiplexed capture (n=14) were mapped to the GRCh38-2020-A-2.0.0 human reference genome using cellranger (v6.0.2) (10X Genomics). Demuxlet^34^ was used to demultiplex each capture into its constituent samples based on genotypes in a common VCF file containing genotype (GT) and genotype likelihood (PL) for each individual. Demuxlet implements a statistical model to determine the likelihood of RNA-seq reads from any given single cell to map to a set of single nucleotide polymorphisms, therefore leveraging natural genetic variation to differentiate between samples from different individuals. Following demultiplexing, the Seurat (v3.2.3 Rv4.1.0) pipeline was used to retain only high-quality cells based on the following criteria: “singlet” as determined by Demuxlet (“doublets” and “ambiguous” cells removed), percent mitochondrial reads < 15%, and RNA read count (nCount_RNA) > 500. Out of the initial 115,698 cells captured across all batches, 92,014 were retained as high-quality singlets.

#### Single-cell ATAC-seq data

FASTQ files from each multiplexed capture (n=14) were mapped to the GRCh38-2020-A-2.0.0 human reference genome using cellranger-atac (v2.0.0) (10X Genomics). Demuxlet^34^ was used to demultiplex each capture into its constituent samples as described above using the same common VCF file. Following demultiplexing, we used the ArchR (v1.0.1, ArchRGenome: hg38) pipeline to filter the data, retaining only high-quality cells. Cell filtering and the creation of ArrowFiles was performed in a single step using the createArrowFiles function on cells with “singlet” demuxlet status and using parameters minTSS = 4 and minFrags = 1000 to further retain only cells with a sufficient signal to background ratio (high accessibility at transcription start sites) and at least 1000 unique nuclear fragments. Across all batches, 58,988 cells were retained as high-quality singlets.

### Clustering, cell type assignments, and UMAP analysis

#### scRNA-seq data

Following quality-control filtering, we split cells first by timepoint giving rise to two groups of cells: Td0 (from D0 samples, n=42,493) and Tm3 (from D90 samples, n=49,521). Since individuals received either the BCG vaccine or placebo, we further split Tm3 cells into two subgroups: BCG (n=37,999) or CTL (n=11,522) – leading to 3 final groups of cells: Td0, Tm3_BCG, and Tm3_CTL. We ran the function SCTransform separately for each group to normalize and scale UMI counts, to identify the most variable features, and to regress out variables corresponding to percent mitochondrial reads or capture. We then integrated the transformed data using the following Seurat functions: SelectIntegrationFeatures (nfeatures=3000), PrepSCTIntegration, FindintegrationAnchors, and IntegrateData. To perform dimensionality reduction downstream of integration we used the functions RunPCA (npcs=30), RunUMAP (dims=1:30), FindNeighbors (dims=1:20), and FindClusters (resolution=0.5). This resulted in 23 preliminary clusters.

To annotate the clusters according to HSPC cell type, we used the FindTransferAnchors function (dims = 1:30, reference.reduction = “pca”, reference.assay = “SCT”, query.assay = “integrated”) to map our integrated scRNA-seq data onto a pre-labelled human bone marrow reference dataset (thawed, stained, sorted, and processed for scRNA-seq as described above) we previously annotated using CellID^35^.

#### scATAC-seq data

Following quality-control filtering and creation of arrow files for each sample, we combined all arrow files into a ArchRProject used in all downstream processing steps. Dimensionality reduction, batch effect correction, clustering, and UMAP visualization were performed using the following functions of the ArchR pipeline: addIterativeLSI (with parameters: iterations = 2, resolution = c(0,2), sampleCells = 10000, n.start = 10, varFeatures = 25000, dimsToUse = 1:30), addHarmony, addClusters (resolution=0.8, reducedDims=Harmony), and addUMAP (nNeighbors = 30, minDist = 0.5, metric = “cosine”). To annotate clusters according to HSPC cell type matching those in the scRNA-seq data, we first performed an unconstrained integration using the addGeneIntegrationMatrix function to broadly map each scATAC cluster to a cell type within our scRNA-seq data. Using this approach, we made the following preliminary assignments:

scATAC clusters “C5”, “C6”, “C19”, “C8” ➔ “HSC”

scATAC clusters “C22”, “C17”, “C2”, “C4”, “C3”, “C1”, “C21”, “C16” ➔ “CMP”

scATAC clusters “C10”, “C23”, “C24” ➔ “GMP”

scATAC clusters “C7”, “C14”, “C15”, “C18” ➔ “MEP”

scATAC clusters “C12”, “C13” ➔ “MLP”

scATAC clusters “C11”, “C9” ➔ “PreBNK”

scATAC cluster “C20” ➔ “unknown”

To generate more detailed cluster mappings (i.e., separating “HSCs” into “HSC c1” or “HSC c2”) we then performed a second-round constrained integration by rerunning addGeneIntegrationMatrix with the newly defined broad group labels, leading to the following final cluster assignments (annotated as: *raw scATAC cluster name/scRNAseq equivalent*):

C1/CMP c1, C2/CMP c1, C3/CMP c2, C4/CMP c1, C5/HSC c2, C6/HSC c1, C7/MEP c1, C8/HSC c2, C9/PreBNK, C10/GMP c2, C11/PreBNK, C12/MLP c2, C13/MLP c2, C14/MEP c1, C15/MEP c3, C16/CMP c1, C17/CMP c3, C18/MEP c3, C19/HSC c2, C20/unknown1, C21/CMP c2, C22/CMP c2, C23/GMP c1, C24/GMP c1

Following exclusion of ‘unknown’ clusters, or clusters for which there were more than 12 samples filtered (see below) the following final cluster names were used downstream of the differential accessibility analysis (below):

C3 -> CMP1, C4 -> CMP2, C5 -> HSC1, C6 -> HSC2, C7 -> MEP1, C9 -> PreBNK, C10 -> GMP1, C12 -> MLP, C14 -> MEP2, C15 -> MEP3, C17 -> CMP3, C18 -> MEP4, C21 -> CMP4, C22 -> CMP5, C23 -> GMP2, C24 -> GMP3

### scATAC-seq Peak calling

We called peaks using the ArchR function addReproduciblePeakSet which utilizes MACS2^36,37^ to call cluster-specific peaks using pseudo-replicates, and then creates a merged peak set using iterative overlap peak merging. For peak calling we used the initial raw, unprocessed alignment data but with added cell type labels derived as described above.

### Pseudobulk estimates

For downstream analyses of scRNA-seq data we summarized single cell expression into pseudobulk estimates for each sample (each unique donor-timepoint pair), allowing a bulk RNAseq-like approach to investigating effects of BCG vaccination on human bone marrow for each cell type. For each of the final 13 unique clusters with a defined cell type label (HSC c1, HSC c2, CMP c1, CMP c2, CMP c3, GMP c1, GMP c2, MEP c1, MEP c2, MEP c3, MLP c1, MLP c2, and PreBNK) we summed raw UMI counts belonging to all cells from the same sample using the sparse_Sums function in textTinyR (v1.1.4). Thus, for each cluster we converted an initial cell by gene (n x m) matrix to a sample by gene (s x m) matrix.

For scATAC-seq data, we summarized single cell peak counts into pseudobulk estimates as described above, only using called peaks instead of genes. As described above, we summed raw peak counts belonging to all cells from the same sample using the sparse_Sums function in textTinyR separately for all clusters (n=24). Thus, for each cluster we converted an initial cell by peak (n x p) matrix to a sample by peak (s x p) matrix.

### Modelling effect of BCG on gene expression and integration with mashr

#### Data filtering/normalization/transformation

Gene expression data: For each cell type, we analyzed pseudobulk gene expression as if it were bulk-RNA sequencing expression data. We first removed any samples for which there were fewer than 20 cells, and any samples for which there was not a matching Td0 or Tm3 timepoint (retaining only paired samples). Lowly expressed genes were filtered by removing all genes for which the median logCPM values calculated for samples in each condition (Td0 CTL, Tm3 CTL, Td0 BCG, Tm3 BCG) were all below 1. Then, we normalized gene expression counts across all samples using the calcNormFactors function implemented in the edgeR R package (version 3.34.1) which utilizes the TMM algorithm (weighted trimmed mean of M-values) to compute normalization factors, and we log-transformed the data using the voom function from the limma package.

Peak accessibility data: Similarly for peak accessibility data, we analyzed each pseudobulk peak count matrix as if it were bulk-ATAC sequencing data. Data was filtered by removing samples with fewer than 20 cells and any samples for which there was not a matching Td0 or Tm3 timepoint. We filtered out low-count peaks for which the median logCPM values calculated for samples in each condition (Td0 CTL, Tm3 CTL, Td0 BCG, Tm3 BCG) were all below a cell-type specific threshold (1.25 for C12 and C15, 1.75 for C9 and C18, 2 for C23, 2.25 for C21, 2.5 for C17, and 1 for all other clusters). Custom thresholds were chosen as we found betas to be positively or negatively skewed when binned by expression level prior to filtering, and that different thresholds for filtering out lowly expressed peaks were required to center these beta distributions. Finally, we normalized peak counts across all samples using the calcNormFactors function in edgeR, and log-transformed the data using the voom function in limma.

#### Model fitting

We wanted to investigate the 90-day impact of BCG vaccination on gene expression and peak accessibility in human bone marrow by comparing expression/accessibility levels from vaccinated individuals at day 90 (after vaccination) and day 0 (prior to vaccination). However, expression and peak accessibility measurements can naturally change across time, independent of whether the individual received the BCG vaccine or only a placebo. Moreover, although individuals assigned to the placebo or BCG cohorts were matched for age, sex, and lack of previous BCG exposure there could be random preexisting baseline differences when comparing the cohorts. To correct for these effects, we independently fit scRNA and scATAC pseudobulk data to a mixed model to estimate the impact of time and cohort assignment on expression/accessibility while also giving an estimate of the independent contribution of BCG-vaccination to changes in expression/accessibility at day 90.

Separately, for each feature (genes or peaks) and each cell type, we fit the following model:

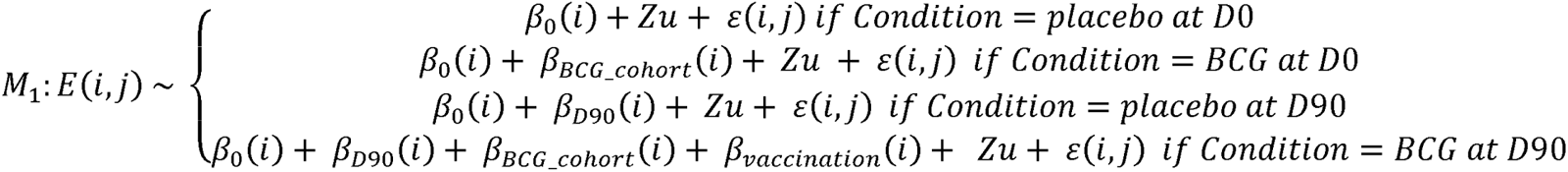

Where *E(i,j)* represents the estimate for each feature *i* and sample *j*. *E(i,j)* is modelled as a function of the fixed effects, β_0_, β_D90_, β_BCG_cohort_, and β_vaccination_, and the random effects Zu. β_0_(i) represents the intercept for the feature i, β_D90_(i) is the natural effect of time on feature *i*, β_BCG_cohort_(i) represents pre-existing baseline differences in feature *i* between the control and BCG cohorts, and β_vaccination_(i) represents the effect of BCG vaccination on feature *i* at D90. The vector *u* is an mx1 vector of random effects to control for individual donor differences where m is the number of unique donors (m=X; m=j/2). Z is an incidence matrix of 1’s and 0’s that maps each sample *j* to one of *m* individuals. The model was fit using the R package EMMREML. *Mashr*

To increase our power to detect BCG-responsive genes shared or unique to each cell type, we applied Multivariate Adaptive Shrinkage in R (mashr version 0.2.57) to outputs from emmreml for scRNA-seq data. We did not apply mashr to scATAC data because peaks accessible enough to pass initial filtering steps are highly cell type specific, decreasing the utility of mashr in this context. For scRNA data, effect sizes were obtained by extracting the betas (β_vaccination_) for each cell type and the standard error of the effect size for each gene was given by taking the square root of varbeta estimates from emmreml. Effect sizes and standard errors for each cell type were arranged into n x m matrices, n being the number of genes and m being the number of cell types. We then fit the mash model using canonical and data driven covariance matrices and then stringently defined significant genes as those with an lfsr < 0.01.

### HSC scRNA-seq subcluster analysis

HSC subcluster analysis was performed by including only cells labelled as HSC c1 and HSC c2 and applying the clustering, pseudobulk, modelling, and mashR steps outlined above.

### MPP score

Calculation of the MPP score was based on known differences in the expression of CD90, CD49f, and CD45RA between MPPs and LT/ST-HSCs. To calculate the score, we first obtained mean expression values for each of the three genes across single cells for each HSC subcluster. The unprocessed mean values were centered and scaled using the *scale* function in R to normalize all values to a mean=0 and standard deviation of 1. Scaled scores for each individual gene were averaged to generate the final composite score.

### Velocyto, Cellrank, and terminal state prediction

We used velocyto^38^ followed by the CellRank^39^ pipeline to determine single cell RNA-velocity measurements and to predict the terminal lineage fate of HSCs from each sample.

We first used the velocyto run10x command to quantify spliced and unspliced read counts (which are required downstream in the pipeline to estimate RNA velocities) for each gene within every cell of our scRNA-seq dataset. Then, separately for cells of each unique donor-timepoint sample, we ran the CellRank^39^ pipeline in python to predict terminal fates of individual HSCs within each sample. Briefly, for each sample, we first removed genes with very low spliced/unspliced mRNA counts, normalized and log-transformed the data, subset on only the top-most variable genes, and computed principal components and moments for velocity estimation using the following CellRank functions: scv.pp.filter_and_normalize (with parameters min_shared_counts=20 and n_top_genes=2000), sc.tl.pca, sc.pp.neighbors (with parameters n_pcs=30 and n_neighbors=30), and scv.pp.moments (with parameters n_pcs=None and n_neighbors=None). Next, we used dynamical modelling to estimate RNA velocities for each single cell using the function scv.tl.recover_dynamics and computed a velocity graph indicating the likelihood that one cell will transition into another based on their RNA velocities and relative positions using scv.tl.velocity(mode=”dynamical”) and scv.tl.velocity_graph. Visualization of these velocity graphs was performed with the function scv.pl.velocity_embedding_stream.

We then used a velocity Kernel to formally predict the terminal lineage fate of each HSC for each sample. We first used the commands VelocityKernel and vk.compute_transition_matrix on the single cell data, pre-processed as described above, to compute a cell-cell transition matrix based on RNA velocity. We combined this velocity kernel with a connectivity kernel to create a less noisy combined kernel (combined_kernel = 0.8 * vk + 0.2 * ck). Using a GPCCA (Generalized Perron Cluster Analysis) estimator, we computed a schur decomposition with g.compute_schur(n_components=20). Finally, we pre-defined all possible terminal states using g.set_terminal_states (with possible states: MLP c1, MLP c2, GMP c1, GMP c2, MEP c1, MEP c2, MEP c3, CMP c1, CMP c2, CMP c3, PreBNK) and then calculated the terminal state probabilities for each HSC using g.compute_absorption_probabilities(use_petsc=True, n_jobs=5, solver=’gmres’).

To compare terminal state differentiation probabilities across time for any given donor we labelled each single HSC with the terminal state towards which it had the greatest differentiation probability. For each donor (excluding donors with fewer than 20 total HSCs) we then computed the percentage of HSCs at day 0 and day 90 having maximal differential probability towards each terminal state and computed the difference across time (%day90 – %day0), leading to a “differentiation-shift” score for each possible terminal state, for each donor. For statistical comparisons of differentiation-shift scores of BCG and placebo groups we used a Wilcoxon test.

### scHINT motif enrichment

#### Preprocessing

Transcription factor motif enrichments and foot printing were performed using HINT-ATAC from the Regulatory Genomics Toolbox. Raw bam files for each 10X capture were split by vaccination cohort, timepoint, and assigned cell type using *samtools* (v1.9) *view*. We focused on comparing BCG samples at D0 and D90, so only BCG samples (i.e., BCG_D0_HSC, BCG_D90_HSC, BCG_D0_CMP, BCG_D90_CMP, …) were processed further. Matching bam files from each capture were merged using *samtools merge* to generate BCG D0 and BCG D90 merged bam files for HSCs, CMPs, GMPs, MEPs, MLPs, and PreBNK (12 total files) for downstream foot printing and motif analyses.

#### Motif enrichment

To determine which motifs were present within DR peaks we performed *rgt-motifanalysis matching* on DR peaks using the JASPAR CORE Vertebrates set of curated position frequency matrices to determine whether specific motifs were significantly enriched we used the *rgt-motifanalysis enrichment* function with cluster-specific DR peaks as the foreground and the shared total peak set as the background for all clusters.

#### Foot printing

To predict the locations of transcription factor footprints we ran the *rgt-hint footprinting* function with parameters --atac-seq --paired-end --organism=hg38 on all peaks for each merged bam file generated in the preprocessing step. To predict which transcription factors were likely bound at each predicted footprint, we used the *rgt-motifanalysis matching* function to find motifs present within footprints.

### Assigning DA peaks to genes and GO enrichment

To investigate which genes were located closest to differentially regulated peaks, we assigned each DR peak to the gene with the closest TSS using the Homer^40^ function *annotatePeaks* with default parameters. This peak-gene association was performed separately for DR peaks within CMPs, GMPs, HSCs, MEPs, MLPs, and PreBNK clusters. To determine whether specific pathways were enriched among genes closest to DA peaks we specified the parameter -GO when running the annotatePeaks function which outputs peak-gene assignments and gene ontology enrichments using DA peaks as foreground peaks and a total peak set (common to all clusters) as background. Output gene ontology enrichment p-values were corrected with the *p.adjust* function in R.

### Regulon analysis

We used pySCENIC^30^, the python implementation of the SCENIC^29^ pipeline, to predict transcription factor activity levels within each cluster. Briefly, we first created a loom file for each cluster for which the analysis was to be performed using the *build_loom* function implemented in the SCopeLoomR package. Then we used the *pyscenic grn* function on the loom object to derive co-expression modules from the single cell expression data. Next, the *pyscenic ctx* function was run with default parameters to search for transcription factor motifs at promoter regions among members of each co-expression module and to trim targets lacking the target transcription factor motifs. Finally, *pyscenic aucell* was used to generate an activity score for each pruned co-expression module for every cell.

To compare transcription factor module activity scores across different conditions, we averaged activity scores for all cells belonging to the same sample to generate average TF activity scores per donor per timepoint. For each donor, we computed the Tm3/Td0 activity score ratio to compute the fold change in activity score across time. Then we compared Tm3/Td0 activity scores between donors of the placebo versus BCG cohorts and used the Wilcoxon rank sum test to derive a p-value.

### Definition of driver TFs

We defined “driver TFs” as transcription factors with evidence of changes in activity score (p < 0.05, see ‘Regulon analysis’ above) and/or TFs whose encoding gene was differentially expressed (lfsr < 0.1) in the scRNA-seq analysis (see above).

### Gene set enrichment analysis

Gene set enrichment analyses (GSEA) were performed using the fgsea R package (version 1.18.0) with parameters: maxSize = 500, nperm=100000. To investigate biological pathway enrichments among BCG-responsive genes, we ordered genes by the rank statistic: – log10(lfsr)*PM where lfsr and PM (posterior mean) were output from running mashr as described above. The rank-ordered gene list was compared with the Hallmark gene sets from the MSigDB collections.

To investigate enrichment of driver TFs (see ‘Driver TFs’ above) among TF motifs found within differentially accessible peaks of downstream progenitors, we ordered motifs by the rank statistic perc*-log10(fdr), where *perc* is the percentage of significant peaks within which the motif is found, and *fdr* is the enrichment of the motif among DA peaks compared to a background of all peaks. For each the progenitor cell type the rank-ordered TF motifs were compared with the list of driver TFs identified for HSCs.

### Elastic net regression

We built an elastic net model using the *glmnet* R package^41^ to determine whether the magnitude of BCG-induced differential accessibility of peaks within progenitors, or differential gene expression in HSCs, was predictive of the log2FC value of cytokine production of PBMCs after BCG vaccination. To choose the optimal value of alpha, we tested alphas ranging from 0 to 1 in increments of 0.1 and chose the alpha that maximized the R2 value between the elastic net predicted IL1B log2FC values, and their experimentally measured values. The regularization parameter lambda was chosen to minimize mean-squared error during n-fold internal cross-validation.

We used a leave-one-out cross-validation approach to generate predicted IL1B log2FC values for each donor. We first separated all samples (each sample corresponding to a donor) into training and test samples and quantile normalized the raw log2FC values (BCG vs. placebo) for each differentially accessible peak, or differentially expressed gene, within each sample to a standard normal distribution. Then, we split the test sample from the training samples, and on the remaining training samples, quantile normalized across samples to a standard normal distribution. For each peak/gene within the test sample, we compared log2FC differential accessibility/expression to the empirical cumulative distribution function for the training samples. This allowed us to estimate the quantile into which the peak/gene fell and to assign this quantile value using the *qnorm* function in R.

### Correlations

All correlations were performed with the *cor.test* function in R with parameter method= “spearman”.

## ACKNOWLEDGEMENTS

We thank the Flow Cytometry Facility and the Genomics Facility at the University of Chicago for their technical support. Computational resources were provided by the University of Chicago Research Computing Center. We thank all members of the Barreiro lab for their insightful comments. This study was supported by an ERC advanced grant (#833247) and a Spinoza grant of the Netherlands Organization for Scientific Research (to M.G.N.). In addition, the work was supported by grant R01-GM134376 to L.B.B., and the UChicago DDRCC, Center for Interdisciplinary Study of Inflammatory Intestinal Disorders (C-IID) (NIDDK P30 DK042086). S.S. is supported by the MSTP training grant T32GM150375.

## AUTHOR CONTRIBUTIONS

L.B.B and M.G.N.conceived the project. L.B.B. directed the study. L.B.B. S.S. M.D. and M.G.N. designed the experiments. C.J.D. L.A.B.J., and M.G.N. collected the samples and designed the original clinical trial. S.S. and A.D. performed the experiments. S.S. led the computational analyses with contributions from R.A.G. and J.S. S.S. and L.B.B. wrote the manuscript, with input from all authors.

