## Supplementary materials for "BCG vaccination impacts the epigenetic landscape of progenitor cells in human bone marrow"

### *Supplementary Material*

#### 1. Supplementary Data

This PDF file includes:

Figures S1 to S4

Table 1-7 legends

#### 2. Supplementary figures

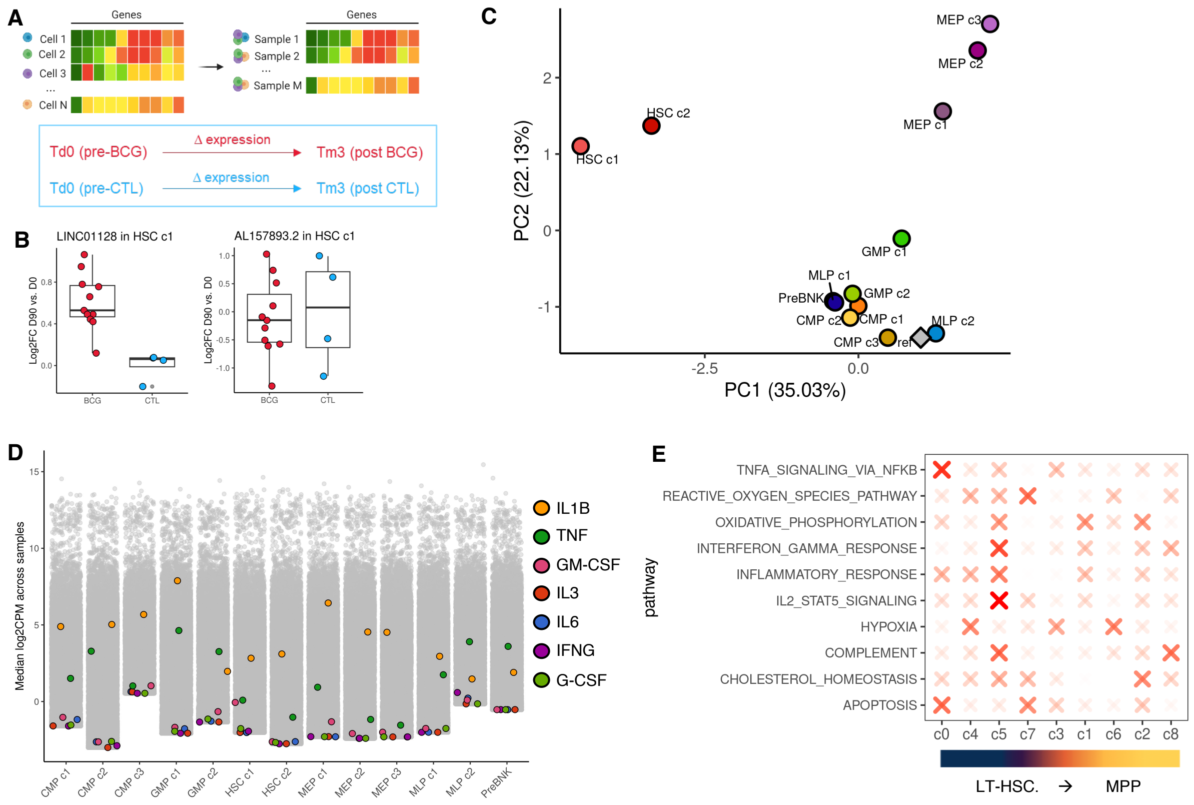


**Supplementary Figure 1. BCG vaccination has heterogenous impacts on gene expression after 90 days. (A)** Schematic showing the general scRNA-seq analysis approach. Raw ‘CELL x GENE’ UMI counts generated through the Seurat pipeline were transformed into ‘SAMPLE x GENE’ pseudobulk matrices for each cell-type/cluster. Pseudobulk expression was fit to a linear model that estimates and corrects for natural expression changes across time in placebo individuals and allows identification of BCG-specific effects on gene expression. **(B)** Example boxplots showing a gene (LINC01128) for which BCG vaccination had a significant differential impact on expression compared to placebo and a non-significant gene (AL157893.2) that exhibited similar across-time changes in expression in both placebo and BCG vaccinated individuals. **C-** Principal component analysis showing cell types clustered by Hallmark pathway enrichment (NES) scores as computed for GSEA in Fig 2C. “Ref” point (gray diamond) is a vector of zeros, representing a baseline unaffected state with no enrichment of any pathway. NES values for pathways with p>0.05 were set to 0. **D-** Raw, normalized log2CPM expression levels of pro-inflammatory and myeloid-differentiation cytokines (IL1B, TNFA, GM-CSF, IL3, IL6, IFNG, G-CSF) in each cluster. **E-** Dot plot of select enriched hallmark pathways within HSC c1 and HSC c2 (Fig 2C) The relative enrichment of the pathway (-log10(padj)) was determined for each HSC subcluster following within-subcluster differential gene expression analysis and GSEA as performed in Fig 2. Enrichment z-score for each pathway was calculated by comparing enrichment scores across subclusters.

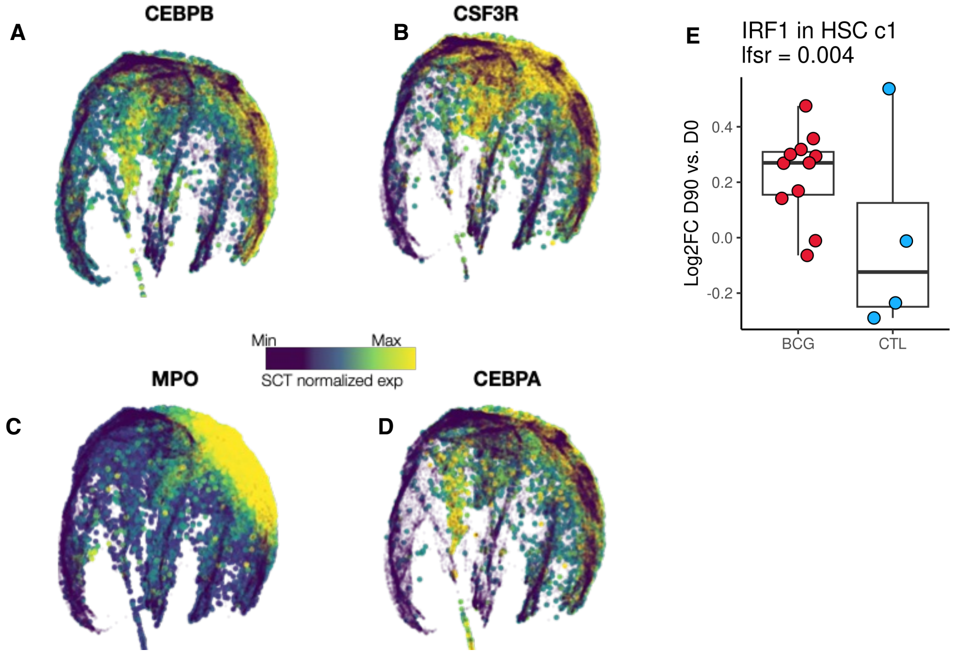


**Supplementary figure 2. The CMP c2 cluster is MPOhi. (A-D)** UMAPs colored by expression levels of monocyte or granulocyte-associated genes (**A.** CEBPB; **B**. CSF3R; **C.** MPO; **D.** CEBPA). **(E)** D90 vs. D0 Log2FC expression of IRF1 (lfsr = 0.004)

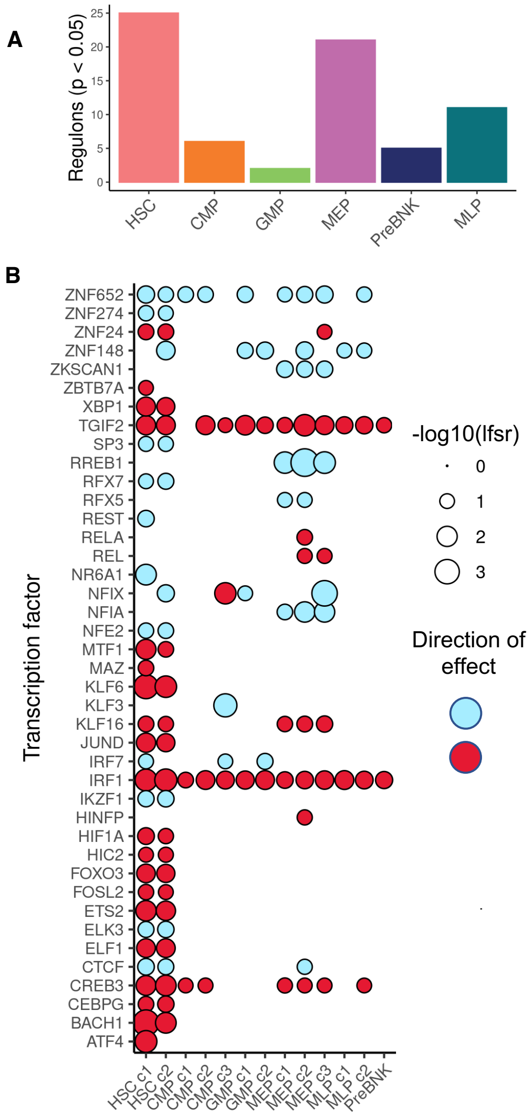


**Supplementary Figure 3. BCG vaccination induces changes in TF activity and gene expression. (A)** Number of transcription factor regulons with differential activity ($p < 0.05$) in each broad cluster group. **(B)** Bubble plot showing TFs that are differentially expressed within each cluster from the RNA-seq data. Red indicates higher expression within BCG vaccinated individuals and blue indicates decreased expression.

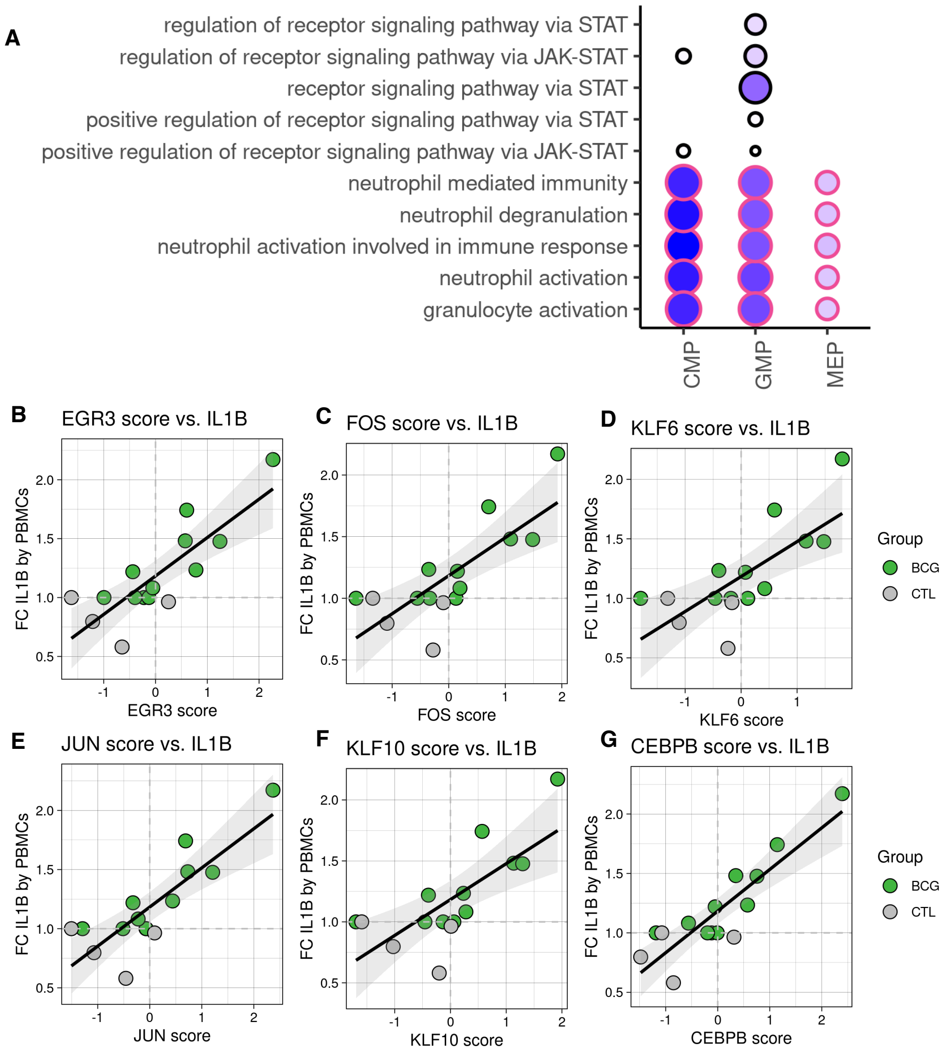


**Supplementary Figure 4. DR peaks in GMPs enrich for innate immune pathways. (A)** DA peaks within each cluster were assigned to the gene with the closest TSS. Gene ontology enrichment analysis was performed for biological process pathways using all peak associated genes as background and genes associated with DR peaks as foreground. Plot circle size and shading darkness are both scaled to –log10(p-value) of enrichment. Pathways related to immunity, immune development, and MAPK signaling are outlined in pink. **(B-G)** Example scatter plots correlating TF activity (regulon) scores in HSCs with fold change IL1B in PBMCs for B) EGR3 (spearman rho = 0.75, p = 0.0013), C) FOS (spearman rho = 0.7, p = 0.0035), D) KLF6 (Spearman Rho = 0.71, p = 0.0033), E) JUN (Spearman Rho = 0.72, p = 0.0023), F) KLF10 (Spearman Rho = 0.74, p = 0.0017), and G) CEBPB (Spearman Rho = 0.78, p = 6e-4).

#### 3. Table legends

**Table 1.** Antibodies and fluorochromes used to stain each bone marrow sample (left), and surface marker combinations used to define each HSPC subtype (right).

**Table 2.** Post-mash lfsr values for genes (includes genes with lfsr<0.1 in at least one cluster) within each cluster.

**Table 3.** Gene set enrichment analysis results for Hallmark pathways within each cluster.

**Table 4.** Summary statistics and coordinate information for DA (differentially accessible) peaks within each cluster.

**Table 5.** Motif enrichment results output by scHINT for each broad cluster group (HSC, CMP, GMP, MEP, MLP, PreBNK)

**Table 6.** Transcription factor activity change (D90-D0) P-values comparing the placebo and BCG vaccination cohorts

**Table 7.** Gene ontology analysis results based on assigning each DA peak to its closest gene, for reactome pathways within each cluster.
